# Mechanisms of damage prevention, signalling, and repair impact disease tolerance

**DOI:** 10.1101/2021.10.03.462916

**Authors:** Arun Prakash, Katy M. Monteith, Pedro F. Vale

## Abstract

The insect gut is frequently exposed to pathogenic threats and must not only clear these potential infections, but also tolerate relatively high microbe loads. In contrast to the mechanisms that eliminate pathogens, we currently know less about the mechanisms of disease tolerance. We investigated how well-described mechanisms that either prevent, signal, control, or repair damage during infection contribute to the phenotype of disease tolerance. We established enteric infections with the bacterial pathogen *Pseudomonas entomophila* in transgenic lines of *Drosophila melanogaster* fruit flies affecting *dcy* (a major component of the peritrophic matrix), *upd3* (a cytokine-like molecule), *irc* (a negative regulator of reactive oxygen species) and *egfr*^*1*^ (epithelial growth factor receptor). Flies lacking *dcy* experienced the highest susceptibility, while loss of function of either *irc* or *upd3* reduced tolerance in both sexes. The disruption of *egfr*^*1*^ resulted in a severe loss in tolerance in male flies but had no substantial effect on the ability of female flies to tolerate *P. entomophila* infection, despite carrying greater microbe loads than males. Together, our findings provide evidence for the role of damage limitation mechanisms in disease tolerance and highlight how sex differences in these mechanisms could generate sexual dimorphism in disease tolerance.

## Introduction

Many insects thrive on decomposing and decaying organic matter containing a diversity of commensal and pathogenic microorganisms. Like most animals, insects have evolved diverse responses to infection, including behavioural avoidance of infection, physical barriers to pathogen entry and a variety of humoral and cellular immune responses [1–3]. These responses have been particularly well described in the fruit fly *Drosophila*, where signalling pathways such as *IMD* and *Toll* are recognised as major contributors to pathogen clearance [1,2,4–7]. In addition to mechanisms that reduce pathogen burdens, it is increasingly recognised that mechanisms promoting disease tolerance are equally important during recovery to a healthy state [8–12]. Disease tolerance is defined as the ability of hosts to maintain health despite harbouring relatively high pathogen loads, a phenotype that has been observed in several species, including insects [13–17] rodents [18,8,19]; birds [20,21]; and humans [22,23,11].

The mechanisms of pathogen clearance are well-described in many animal species [24], but we currently know less about the mechanisms underlying disease tolerance. Given that tolerance reflects the ability to maintain health independently of pathogen clearance, we might expect tolerance mechanisms to be related to processes such as detoxification, reduction of inflammation, or tissue damage control and cellular renewal [9,11,25]. Genome-wide association or transcriptomic studies in *Drosophila* have highlighted potential candidate genes underlying phenotypic variation in disease tolerance [14], but it remains unclear how many of these genes interact with known mechanisms of immunity and recovery.

Furthermore, almost all candidate genes for disease tolerance in *Drosophila* arise from systemic infections [14,26,27], leaving a gap in our knowledge about disease tolerance during orally acquired infections, which are especially relevant in the context of the ecology of most insects, that thrive on decomposing and decaying organic matter containing a diversity of both commensal and pathogenic microorganisms [28–30]. The insect gut is therefore frequently exposed to pathogenic threats and must be able not only to detect and clear these potential infections, but also be able to repair the resulting damage to gut tissues in order to tolerate relatively high numbers of ingested pathogens.

Here, we aimed to specifically test how well-established mechanisms that either prevent, reduce, or repair tissue damage contribute to the phenotype of disease tolerance. The *Drosophila* gut is a compartmentalized tubular organ which is structurally and functionally similar to the vertebrate intestinal tract [2,31–33]. We can consider several stages comprising gut defence in *Drosophila (****Fig***. *SI-1)*. The first involves the physical barrier of the gut epithelia and the peritrophic matrix (*PM*), which is a layer of chitin and glycoproteins that lines the insect midgut lumen. The *PM* is functionally analogous to mammalian mucus membrane in the digestive tract and acts as the first line of defence against invading pathogens [2,34]. A major component of the *PM* is *drosocrystallin* (*dcy*). Loss-of-function mutations in *dcy* increase the permeability of the peritrophic matrix to larger molecules and allow leakage of microbial cells, including pathogens, into the haemolymph. *Dcy* deficient flies therefore exhibit increased susceptibility to oral bacterial infections [2].

Another mode of defence during gut infections is the production of reactive oxygen species (*ROS*) by the gut epithelia. For example, in response to ingested *Pseudomonas entomophila, ROS* production is induced by two NADPH enzymes-*nox* (NADPH oxidase) and *duox* (dual oxidase), while *irc* (immune-reactive catalase) negatively regulates *ROS* production once the infection threat is controlled, which otherwise, would lead to cytotoxic effects [2,35,36]. *ROS* production not only targets pathogens directly, but also plays additional roles in triggering signalling pathways that lead to the production of *IMD-* or *Toll*-responsive antimicrobial peptides [36–39].

The final stage in gut defence is to repair the damage caused during the infection. Damage-signalling cytokine-like molecules *upd3* are released from damaged cells which trigger the *Jak/Stat-*pathway, stimulating the proliferation of intestinal stem cells (*ISCs*) and their differentiation into enterocytes (*ECs*) via *egfr*^*1*^ (epidermal growth factor receptor) signalling [40–42]. Flies lacking *Jak/Stat* or *Egfr* are therefore highly susceptible to bacterial infections due to their inability to repair and renew damaged tissue [40,42,43].

To investigate how these mechanisms of damage prevention (*dcy*), signalling (*upd3*) control (*irc*) and renewal (*egfr*) contribute to disease tolerance during gut infections we employed oral infections in *Drosophila* transgenic lines with loss-of-function in each of these genes on a common genetic background (*w*^*1118*^). We orally challenged these flies with three infection doses of *Pseudomonas entomophila* and then quantified their effects on survival, pathogen loads and disease tolerance responses during period of peak infection burden.

## Materials and methods

### Fly strains

The following fly stocks were obtained from the Bloomington Stock Centre, Indiana: *dcy* (*w*^*1118*^; Mi{ET1}^CrysMB08319^; FB*#26106*) [44], *irc* (*w*^*1118*^; Mi{ET1}^IrcMB11278^; *FB#29191*) [45], (*#2079*), *upd3* (*w*^*1118;*^ P{XP}^upd3d11639^; *FB#19355*) [46]. These lines were subsequently isogenised by backcrossing onto the same *w*^*1118*^ background (*VDRC stock# 60000*) for at least 10 generations. The egfr^t1^ mutant was a kind gift from Carla Saleh (Pasteur Institute, Paris) and previously isogenised to *w*^*1118*^ first by replacing the chromosomes not containing the mutation using balancer chromosomes and then by backcrossing at least ten times to *w*^*1118*^ line [47]. All fly lines were maintained in plastic vials (12 ml) on a standard sugar-cornmeal medium at a constant temperature of 25°C (±2°C).

### Bacterial culture preparation

To test the impact of bacterial infection on fly survival we used the gram-negative bacteria *Pseudomonas entomophila*, that commonly infects a broad range of insects and other invertebrates. In flies, *P. entomophila* infection mainly occurs in the intestinal epithelium and eventually causes death [35]. To obtain bacterial cultures for oral exposure, we inoculated frozen isogenic bacterial stock cultures stored at −80°C onto fresh 15ml LB broth (media composition) and incubated overnight at 37°C with shaking at 120rpm (revolutions per minute). The overnight cultures were diluted 1:100 into 500 ml of fresh LB broth and incubated again at 30°C with shaking at 120rpm. At the mid-log phase (OD_600_=0.75), we harvested the bacterial cells by centrifugation at 5000rpm for 15 min and re-suspended the bacterial pellet in 5% sucrose [48]. The final inoculum was adjusted to three different bacterial concentrations or infection dose OD_600_=10 (low dose), OD_600_=25 (medium dose) and OD_600_=45 (high dose).

### Experimental design

Measuring tolerance as a linear reaction norm requires collecting matching data on survival and pathogen loads, ideally from the same individual. However, this is challenging in the fly model because quantifying microbe loads requires destructive sampling. Instead, we considered the vial as the unit of replication, and employed a split vial design (***Fig***. *SI*-2). In total, we set up 500 infection vials, split across 2 experimental blocks (n=10 vials for OD_600_=10 and OD_600_=45; n=30 vials for OD_600_=25, for each combination of sex (2) /fly line (5) – with each vial containing 27-30 flies for all doses). Following oral bacterial exposure (see below) each vial containing 25 flies of each infection treatment, sex and fly line combination were split into two vials for measuring (1) survival following infection (15 flies/combination) and (2) internal bacterial load (10 flies/combination) (***Fig***. *SI-2)*. This split-vial design allowed us to use replicate-matched data for both the proportion of flies surviving and the average bacterial load for each replicate vial to estimate the linear relationship between fly survival and internal bacterial load for each fly line.

### Oral infection assay

Before infecting flies we prepared infection vials by pipetting 350µl of standard agar (1L triple distilled H_2_O, 20g agar, 84g brown sugar, 7ml Tegosept anti-fungal agent) onto the lids of 7ml tubes (bijou vials) and allowed it to dry. Simultaneously, we starved the experimental flies in 12ml agar vials for 4-5 hours. Once the agar in the bijou lids dried, we placed a filter disc (Whattmann-10) in the lid and pipetted 80µl of bacterial culture directly onto the filter disc. For control (mock) infections, we replaced bacterial culture with a 5% sucrose solution. We then orally exposed flies inside the bijou vials for 18-hours and then transferred the flies onto fresh vials containing standard sugar-cornmeal medium [48].

### Bacterial load measurement

To test whether variation in mortality of experimental flies after *P. entomophila* infection is explained by the ability to clear infection, we measured bacterial load using 3–5-day old flies (w^1118^ and transgenic flies) following three different doses of *P. entomophila* enteric infection. We used either OD_600_=10 (low dose), OD_600_=25 (medium dose) or OD_600_=45 (high dose). For OD_600_=25 (medium dose), we measured bacterial load at three timepoints (immediately after oral exposure 0-15 minutes, 24-hours, and 96-hours) following *P. entomophila* infection. To confirm oral bacterial infection, we thoroughly surface-sterilised flies (group of 3) with 70% ethanol for 30-60 seconds and then rinsed twice with sterile distilled water. We plated the second wash on LB agar plates and incubated overnight at 30°C to confirm that surface bacteria were successfully removed after alcohol sterilization. We transferred flies onto 1.5ml micro centrifuge tubes and homogenized using a motorized pestle for approximately 30-60 seconds in 100µl LB broth (n=30 homogenates/sex/infection treatment/fly line). We performed serial dilution of each homogenate up to 10^−6^-fold and added 4μl aliquot on a LB agar plate. After this, we incubated the plate overnight for 18-hours at 30°C and counted the resultant bacterial colonies manually. We note that mock-infected control fly homogenates did not produce any colonies on LB agar plates [48].

### Statistics

#### Survival following oral infection

We analysed survival data using a mixed effects Cox model using the R package ‘coxme’ [49]. We specified the model as: survival ∼ fly line * treatment * sex * (1|vials/block), with ‘fly line’, ‘treatment’ and ‘sex’ and their interactions as fixed effects, and ‘vials’ nested in ‘block’ as a random effect for *w*^*1118*^ and flies deficient of damage prevention and repair mechanisms in gut-epithelia.

#### Internal bacterial load

We found that residuals of bacterial load data were non-normally distributed when tested using Shapiro–Wilks’s test. Hence, we first log-transformed the data and then confirmed that the log-transformed residuals were still non-normally distributed. Subsequently, we used a non-parametric Kruskal-Wallis test to test the effects of each fly line, that is, control *w*^*1118*^ and transgenic flies for males and females separately following oral *P. entomophila* infection.

#### Measuring disease tolerance

Finally, to understand how damage signalling and repair mechanisms affect disease tolerance in males and females during oral *P. entomophila* infection, we analysed the linear relationship between fly survival against bacterial load by fitting linear models [8,50,10,16,51,15]. We assessed differences in disease tolerance (fly survival with increasing bacterial load) by fitting ‘fly line’ and ‘sex’ as categorical fixed effects, ‘average bacterial load (log_10_)’ as a continuous covariate, and their interactions as fixed effects. Significant interaction effects between fly line and bacterial load would indicate that the slope of the relationship between fly survival and load varies between fly lines, that is, the tolerance response differs between lines. Because our interest was to quantify the effect of damage prevention and repair mechanisms on disease tolerance, we compared the slope estimates of each of the transgenic lines with the slope of *w*^*1118*^ line using a pairwise comparison (f-test).

## Results

### 1. Flies lacking *dcy* are more susceptible to oral *P*. *entomophila* infections than those lacking components that minimise, signal or repair damage

Following oral infection with three different doses of *Pseudomonas entomophila*, flies with disrupted components of damage prevention (*dcy)*, signalling (*upd3)*, renewal (*egfr*^*1*^)and regulation (*irc)*, were all significantly more susceptible to oral *P. entomophila* infections compared to *w*^*1118*^ flies (*Fig. 1, Table SI-1*; *Fig. 1A* for infection dose *OD*_*600*_*=25; Fig. 1B* for infection dose *OD*_*600*_*=10; Fig. 1C* for infection dose *OD*_*600*_*=45*). Among these lines, *dcy* knockouts were particularly susceptible to infection *Fig.1* and *Fig. SI-3* for survival, *Fig. 1, Table SI-2* for hazard ratio. The effect of each gene disruption on the survival of flies following infection was similar in males and females (fly line x sex x treatment interaction = non-significant, *Table SI-1*), compared to control w^1118^ flies at both lower (*OD*_*600*_*=10*) and higher infection (*OD*_*600*_*=45*) doses (*Fig. 1, Table SI-1*).

**Figure 1.**
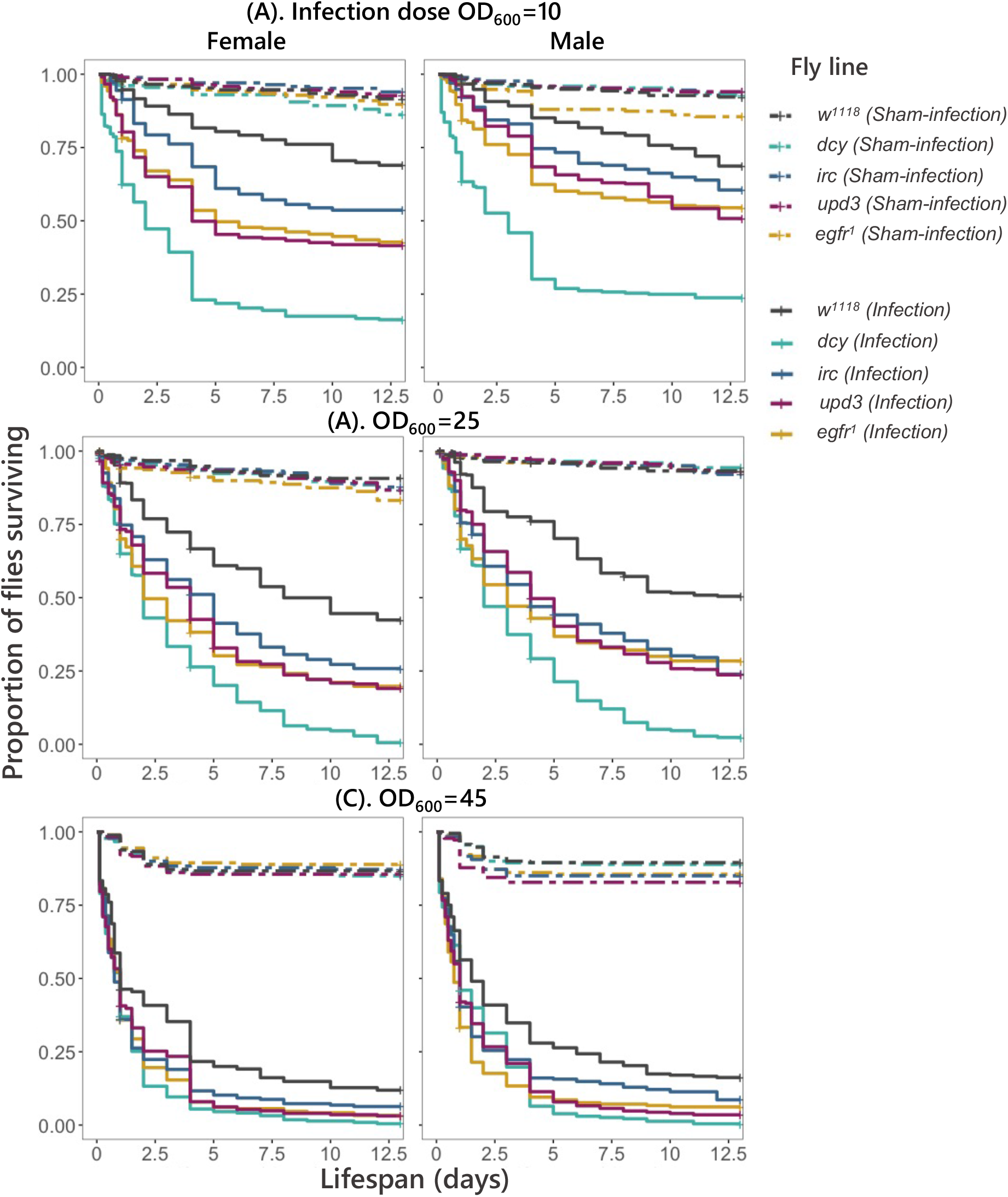
Kaplan Meier curves for females and males of *w*^*1118*^ and transgenic flies with loss-of-function in gut-epithelial response, exposed to oral *P. entomophila* of infection dose ***(A)***. *OD*_*600*_*=10*, n=10 vials per treatment/sex/fly line **(*B*)**. *OD*_*600*_*=25*, n=30 vials per reatment/sex/fly line ***(C)***. *OD*_*600*_*=45*. n=10 vials per treatment/sex/fly line. For all doses, 15-17 flies per replicate vial).

### 2. Both w^1118^ and flies with disrupted tissue damage prevention and repair mechanisms show sex differences in bacterial load during oral infections

The higher susceptibility of all transgenic flies to oral bacterial exposure could either be caused by their inability to supress the bacterial growth or due to their inability to tolerate the damage inflicted during oral infection. To distinguish between these mechanisms, following the overnight oral exposure to *P. entomophila*, we first quantified internal bacterial loads at 15 minutes, 24-hours and 96-hours. We observed a peak in microbe loads at 24h post infection in all fly lines (see ***Fig***. *SI-4, Table SI-3)*. All fly lines showed sex-differences at 24-hours peak load, though by 96-hours following oral infection this sex difference was no longer present in flies lacking *upd3* or *egfr* expression (***Fig***. *SI-4*, ***Table*** *SI-3*). Flies lacking *irc* expression always exhibited levels of bacterial load similar to the w^1118^ flies across all infection doses (***Fig***. *2*, ***Table*** *SI-3*).

**Figure 2.**
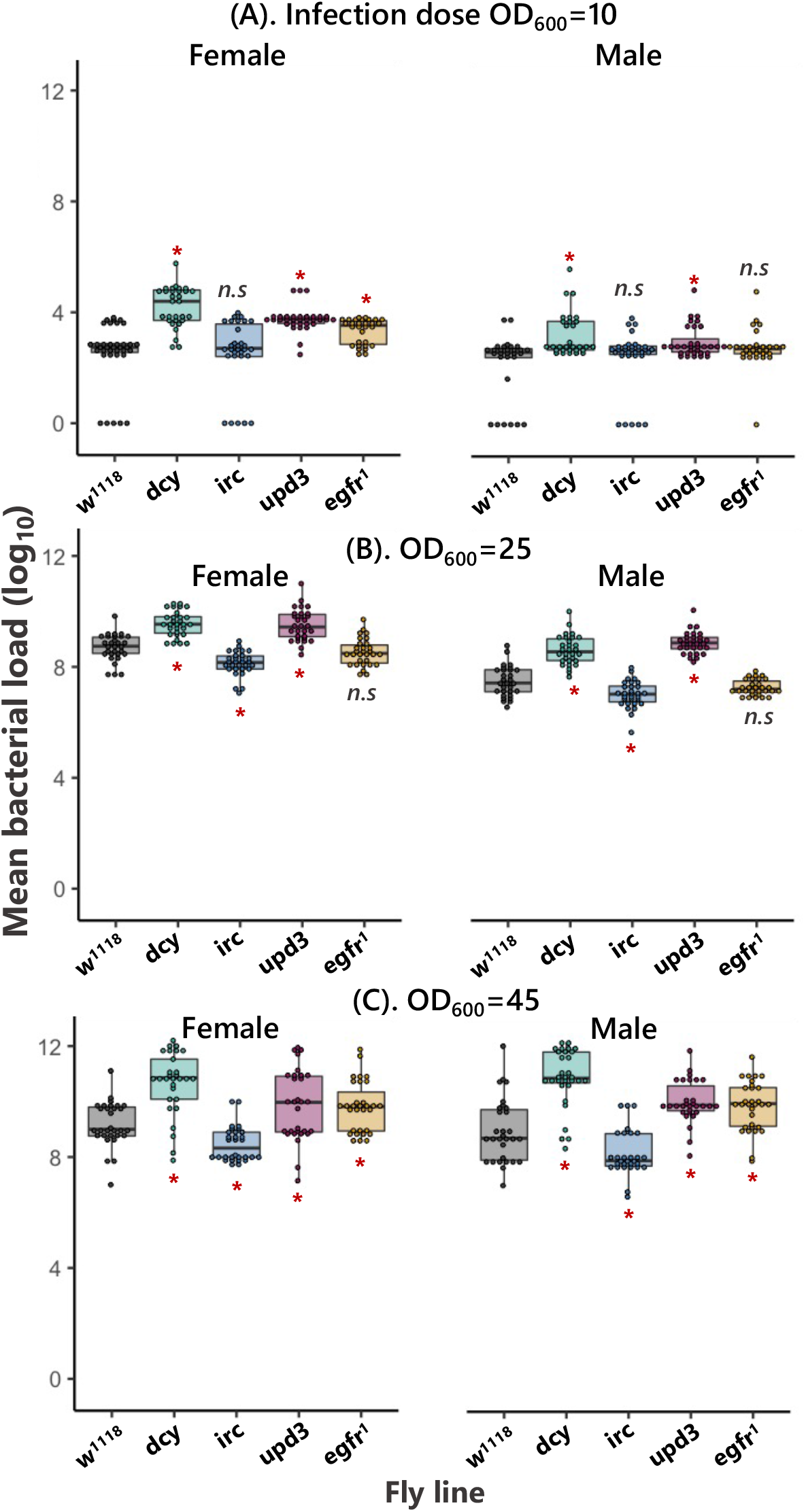
Bacterial load measured as CFUs-colony forming units using infection dose ***(A)***. *OD*_*600*_*= 10 (low dose)* ***(B)***. *OD*_*600*_*=25 (medium dose)* and ***(C)***. *OD*_*600*_*=45 (high dose)* after 24-hours following oral bacterial exposure with *P. entomophila* for *w*^*1118*^ flies and transgenic lines [n=30 vials of which 12-15 flies/treatment/sex/fly line for bacterial load measurement]. Significantly different transgenic lines from control w^1118^ flies are denoted with ‘*’ analysed using pairwise comparisons between w^1118^ and for males and females respectively.

### 3. Damage repair mechanisms mediate sex differences in disease tolerance during oral bacterial *P. entomophila* infections

While some of the variation in survival between fly lines (***Fig***. *1*) may be explained by variation in resistance [that is, their ability to clear infection (***Fig***. *2*)], some of that variation may also arise due to differences in tolerance. We were therefore interested in measuring disease tolerance in these lines using the reaction norm of survival relative to bacterial loads, where the slope of the linear relationship reflects the degree of tolerance: steep negative slopes indicate a rapid mortality with increases in pathogen loads (low tolerance), while less steep or flat slopes reflect relatively more tolerant host [8,50,16,52]. We focused on flies infected with the intermediate dose (*OD*_*600*_*=25*), as we had 30 replicate-paired measurements of survival and microbe loads. We found that the disruption of damage prevention, signalling and repair genes showed reduced tolerance, measured as the rate at which fly survival changed with bacterial load relative to *w*^*1118*^ flies (***Fig***. *3*, ***Table*** *2*). Both males and females lacking the major component of the peritrophic matrix *dcy* showed significantly reduced survival (***Figs***. *1* *and 3*), indicating the general importance of damage preventing epithelial barrier in defence against oral infections [2]. Flies lacking the damage renewal mechanisms *egfr*^*1*^ showed sex-differences in disease tolerance (***Fig***. *3*, ***Table*** *1* *and 2*). We found that the disruption of *egfr*^*1*^ resulted in males, but not females becoming less tolerant of *P.entomophila*, showing a much steeper decline in survival with increasing microbe loads compared to females (**Fig**. 3). Notably, egfr^1^ males are less tolerant than w1118 despite harbouring comparable microbe loads (Fig. 2).

**Table 1.** Summary of ANCOVA. To assess differences in infection tolerance (fly survival with increasing bacterial burden) following oral *P. entomophila* infection with *OD*_*600*_*=25* infection dose, after 24-hours. We analysed ANCOVA and fitted ‘sex’ as categorical fixed effects, ‘average bacterial load’ as a continuous covariate and their interactions as fixed effects for each of the fly lines (*w*^*1118*^ and flies lacking damage prevention and repair mechanisms in gut-epithelia).

| **Sex** | **Source** | **df** | **Sum of Sq.** | **F ratio** | **p** |
| --- | --- | --- | --- | --- | --- |
| *Female* | Fly line | 4 | 66.33 | 71.6 | <0.01 |
|  | Bact. load | 1 | 19.27 | 83.3 | <0.01 |
|  | Fly line x Bact. load | 4 | 10.02 | 10.8 | <0.01 |
| *Male* | Fly line | 4 | 84.60 | 62.8 | <0.01 |
|  | Bact. load | 1 | 28.39 | 84.3 | <0.01 |
|  | Fly line x Bact. load | 4 | 14.88 | 11.0 | <0.01 |

**Table 2.**
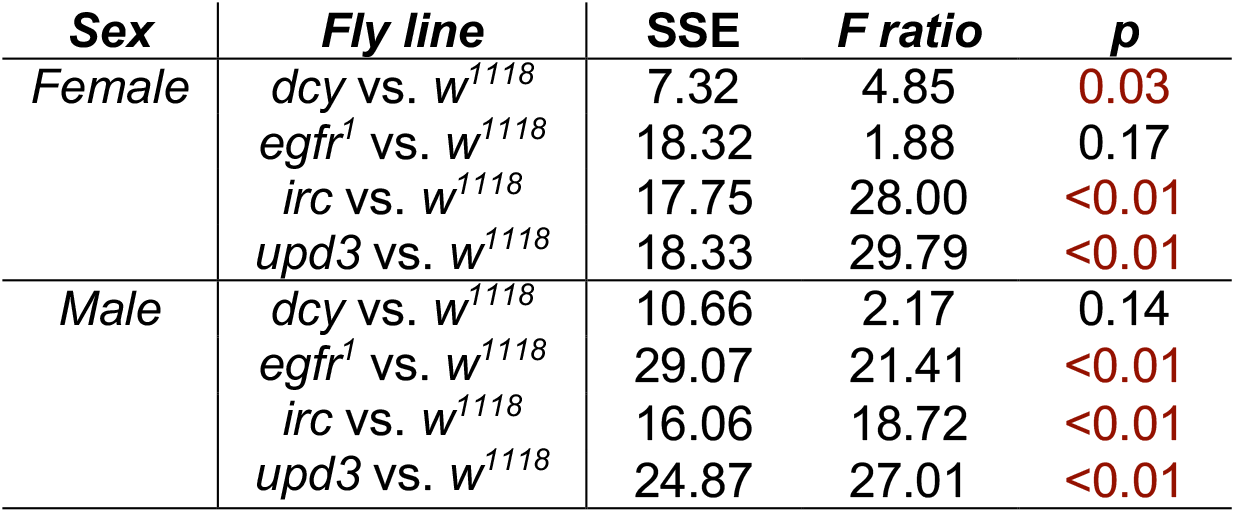
Summary of pairwise comparisons (f-test) of linear slope estimates from linear reaction norm for *w*^*1118*^ flies and flies lacking damage prevention and repair mechanisms in gut-epithelia.

**Figure 3.**
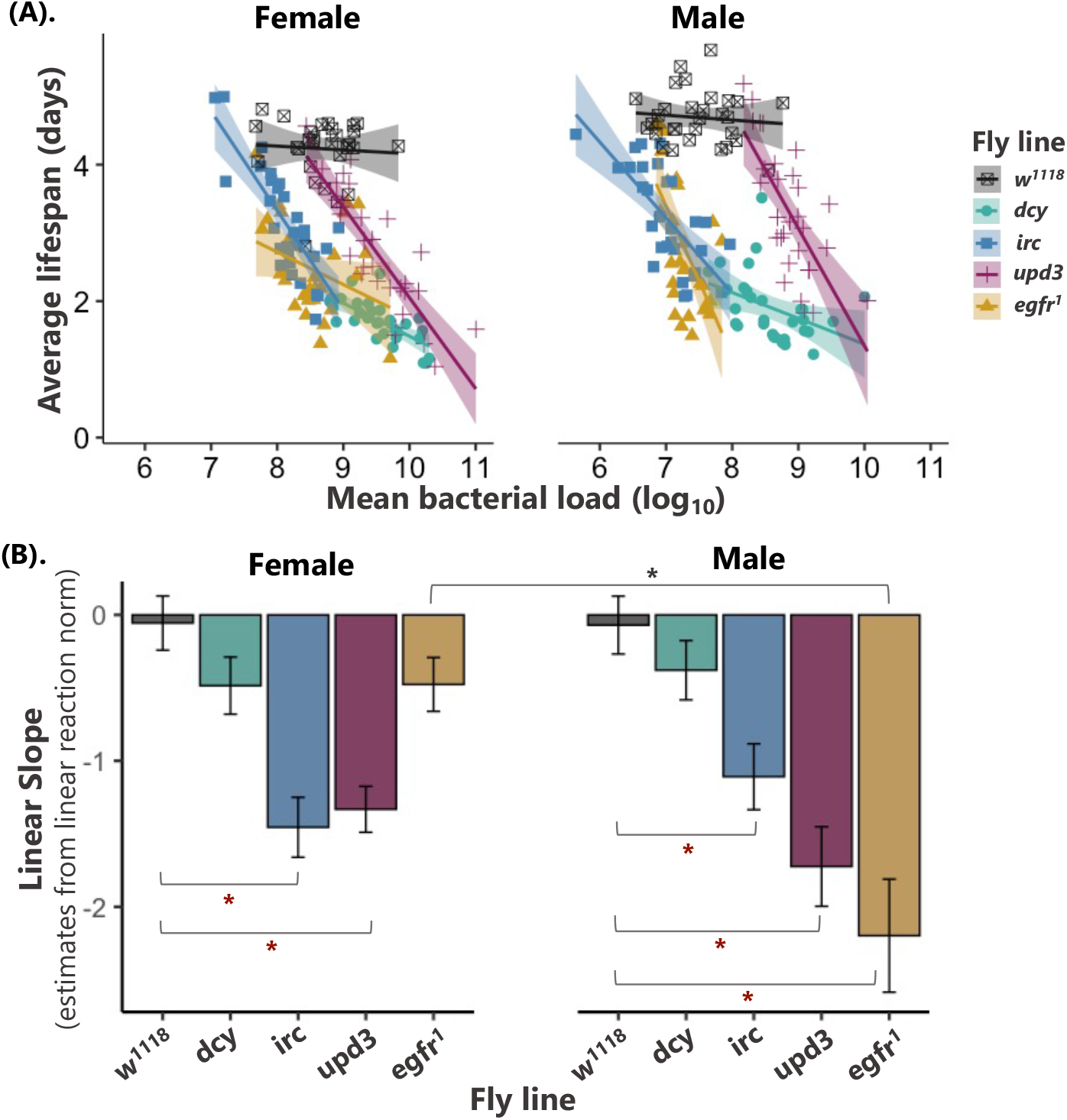
**(A)**. The relationship between fly survival (measured as average lifespan) and bacterial load (as mean CFUs – colony forming units), analysed using linear models for female and male flies (*w*^*1118*^ and flies lacking damage prevention and repair mechanisms in gut-epithelia). Each point shows data for average lifespan and mean bacterial load (CFUs) of 30 vials (with each vial containing 25 individual flies/fly line/sex combination) after 24-hours post oral bacterial exposure. The data shown here are for the medium infection dose *OD*_*600*_*=25*. **(B)**. shows the slope of the linear reaction norm extracted from the linear models. [grey ‘*’ indicates significant difference in tolerance between males and females in tolerance to bacterial infections (interaction between the bacterial load and the sex for each fly line measured using ANCOVA, Table 1), maroon ‘*’ indicates that fly lines (flies lacking damage prevention and repair mechanisms in gut-epithelia) are significantly different from *w*^*1118*^, analysed using pairwise f-test from linear norm estimates see Table 2].

## Discussion

In the present work we tested how mechanisms of damage prevention (*dcy*), signalling (*upd3*) control (*irc*) and renewal (*egfr*) contribute to disease tolerance during enteric infection. We present evidence that all these mechanisms contribute to disease tolerance during bacterial gut infection, and that some of these effects are sexually dimorphic. Previous transcriptomic, genome-wide association (GWAS) or microarrays studies have identified several candidate genes associated with disease tolerance, including - *CrebA, grainyhead and debris buster, dFOXO* [14,26,27,53]. However, all this work has focused on flies infected systemically by directly injecting bacteria into the fly. Here, we investigated tolerance during the natural oral route of infection, and we took a more targeted approach to specifically investigate how some of the well-described tissue damage prevention and repair mechanisms affect disease tolerance during enteric bacterial infections.

Though repairing infection-damage is crucial to fly survival, we found that flies lacking damage-preventing (*dcy*) are particularly susceptible to oral infections compared to those lacking components that minimise, signal or repair damage. In other words, preventing damage is clearly preferable to repairing damage from the perspective of fly survival. This result is consistent with previous work showing that loss-of-function in *dcy* increases the peritrophic matrix width making the gut leaky and compromising gut barrier function during oral bacterial infections [54,44,55,56,2]. We also found increased bacterial loads relative to the w^1118^ control line, measured after 24-hours following infection in both male and female *dcy* knockouts. This is likely because of the combination of leaky gut and pore-forming toxin produced by *P. entomophila* [2] resulting in higher bacterial growth in the fly haemolymph.

In the case of *upd3*-knockout flies, we found reduced survival and higher bacterial loads compared to *w*^*1118*^ flies. Previous work has shown that in response to *P. entomophila* infections, excessive reactive oxygen species (*ROS*) produced by host cells destroy the gut epithelia and block the gut repair process [57,58,55]. The *JNK* and *Hippo* pathways are activated in damaged enterocytes (*ECs*), which produce *upd3*, in turn activating the *Jak/Stat* pathway in intestinal stem cells (*ISCs*). We found that both male and female *upd3* knockout flies showed reduced tolerance and this is likely because in the absence of *upd*3 released from damaged cells the *Jak/Stat*-pathway activation is reduced, which is further necessary for *ISCs* proliferation and differentiation into enterocytes (*ECs*), together renew the damaged tissues [55,2,31]. We also found that functional disruption of *irc* results in lower bacterial loads, and this is expected because loss of this negative regulator of ROS results in higher *ROS* levels [2], leading to improved bacterial clearance.

Regarding the effects of these damage limitation mechanisms on disease tolerance, overall, we found that both male and female *w*^*1118*^ flies were quite tolerant of enteric bacterial infections (reflected in their relatively flat tolerance slopes, ***Fig*** *3*. ***Table*** *2*), while disrupting most damage prevention and repair mechanism lowered disease tolerance (decline in slopes relative to w^1118^). While we found reduced tolerance in all knockout lines, disrupting some components of damage limitation had particularly severe effects on disease tolerance. Significant reductions in disease tolerance were observed in flies with disrupted *irc* and *upd3*, and in these fly lines the effect was comparable in both sexes. *Irc*-deficient flies are unable to regulate *ROS* levels which would lead to increased cytotoxic effects [2,36] while *upd3* cytokine molecules are important for the activation of the *Jak/Stat* pathway [43,40]. In case of *dcy*-knockout flies, health did not deteriorate much further, possibly because it was already too poor to worsen further (***Fig***. 3).

We observed the fastest decline in tolerance in male flies lacking *egfr*^*1*^, but the disease tolerance of female *egfr*^*1*^ knockouts appeared unaffected. This sex difference in tolerance may arise as the result of sex differences in gut physiology and repair. Recent work has demonstrated that during oral *Ecc15* infection, males showed significantly lower gut *ISCs* in response to infection, while female had higher *ISCs* and were resistant to infection and other stress [59]. The differentiation and proliferation of *ISCs* (intestinal stem cells) via *Jak/Stat* signalling into *ECs* (enterocytes) via *egfr* is indispensable for tissue damage renewal. Loss of *egfr*^*1*^ signalling might therefore be felt more severely in males than in females, explaining why male but not female *egfr*^*1*^ knockouts showed severe a decrease in disease tolerance. To date, only a small proportion of studies have compared sex-differences in intestinal immunity, with the majority of work focusing on one particular sex, usually females [60,41,59,61].

Another possibility for the observed sex-difference in damage repair process, might relate to gut-plasticity such as gut-remodelling. For instance, females of mammals such as mice extensively remodel their guts, increasing both digestive and absorptive capacity depending on the nutritional demands of lactation [62]. The remodelling of the gut might be one of the possible driving factors for dimorphism in gut immunity, since males and females differ in their nutritional needs [61]. Studies using *Drosophila* have shown that males and females can make different diet or nutritional choices in accordance with their reproduction role and demand [63] and the *Drosophila* midgut plastically resizes in response to changes in dietary sugar and yeast [64]. Whether gut remodelling and nutritional choice-demand causes sex-differences in damage repair process during disease tolerance remains a question for future research.

### Concluding remarks

Although host mechanisms of immune-mediated clearance are key for pathogen defence and elimination, there is an increasing appreciation that additional defence mechanisms which prevent, signal, repair or renew the extent of tissue damage are also key to infection outcomes by promoting disease tolerance [25,65]. Tissue damage repair mechanisms that promote disease tolerance are interesting from a therapeutic perspective [9,10,66]. For instance, in mice, mechanisms that prevent or repair damage have been shown to confer disease tolerance during malarial *Plasmodium* infection and also during co-infections by pneumonia causing bacteria (*Legionella pneumophila*) and influenza virus [67,68]. Understanding how tissue damage prevention and repair mechanisms contribute to disease tolerance may also help explain how other arthropods are able to vector bacterial and viral infections without substantial health loss [15,69,70]. In summary, our results show that the disruption of tissue damage repair processes resulted in severe loss of disease tolerance and highlight how sex differences in some damage repair mechanisms could generate sexual dimorphism in gut immunity.

## Data accessibility

All the raw data and analysis code can be accessed under a Creative Commons 4.0 License at https://doi.org/10.5281/zenodo.6510215 [71]. An earlier preprint is available from the biological preprint server *bioRxiv* at: https://www.biorxiv.org/content/10.1101/2021.10.03.462916v1 [72].

## Acknowledgements

We thank V. Mongelli and CM. Saleh for sharing the *egfr* line and FM. Waldron for backcrossing *irc, upd3, dcy* fly lines in *w*^*1118*^ background. V. Doublet and E. Robertshaw for laboratory help. JC. Regan, AB. Pedersen, DH. Nussey, DJ. Obbard, TS. Salminen and Ashworth fly group members for helpful discussion. Finally, we thank A. Reid, A. Fulton, L. Rowe and J. King for help with media preparation. *Figs. SI-1 and SI-2* were created using Biorender.

## Authors’ contribution

A.P and P.F.V conceived and designed the experiments; A.P carried out experiments with assistance from K.M.M; A.P analysed the data; A.P and P.F.V wrote the manuscript.

## Supplementary information

### Supplementary figures

**<u>Figure SI-1</u>**. Schematic representation of *Drosophila* gut-epithelial immune response following enteric infection.

**Figure SI-2**. Split-vial experimental design.

**<u>Figure SI-3</u>**. Estimated hazard ratios

**Figure SI-4**. Bacterial load measured at 24 hours after OD_600_=25 of oral *Pseudomonas entomophila* infection

### Supplementary tables

**Table SI-1**. Summary of mixed effects Cox model,

**Table SI-2**. Summary of estimated hazard ratio relative to w^1118^ from the cox proportional model.

**Table SI-3**. Summary of log_10_ transformed bacterial load data after exposure to 3 different concentration (doses) *P. entomophila*

### Supplementary figures

**Figure S1.**
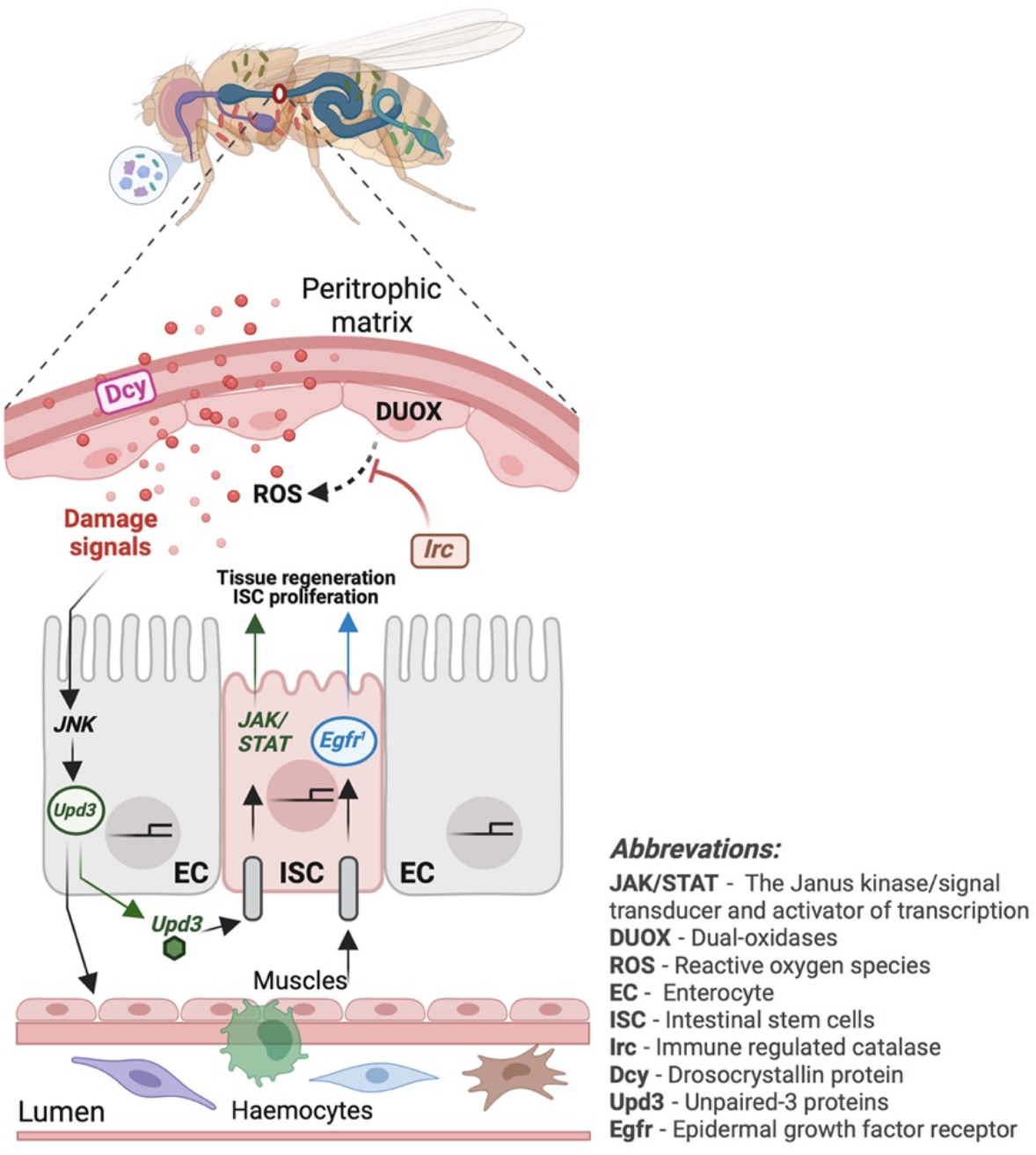
Schematic representation of *Drosophila* gut-epithelial immune response following enteric infection. Several stages of damage prevention and repair, epithelial response in the *Drosophila* gut can be found. ***Stage-I***: involves the physical barrier the peritrophic 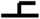 matrix (*PM* - layer of chitin and glycoproteins) that lines the *Drosophila* midgut lumen. A major component of the *PM* is ***dcy*** (*drosocrystallin*) and loss-of-function mutations in *dcy* increase the width of the *PM*, increasing permeability to pathogens. ***II***: production of reactive oxygen species (*ROS*) by the gut epithelia - enteric infections cause massive destruction of the gut epithelium and in response to ingested pathogens, *ROS* production is induced by *nox* (NADPH oxidase) and *duox* (dual oxidase), while ***irc*** (immune-reactive catalase) negatively regulates *ROS* production by suppressing the cytotoxic effects of *ROS* once the infection threat is lessened, avoiding immunopathology. In addition, *ROS* production is known to trigger signalling pathways the *IMD* or *Toll* that leads to the production of antimicrobial peptides (*AMPs*) critical for pathogen clearance in the gut. ***III***: damage renewal process after infection - The final stage in gut defence is to repair the damage caused during enteric infections. Damage-signalling cytokine-like molecules ***upd3***, released from damaged cells trigger the *Jak/Stat*-pathway, stimulating the proliferation of intestinal stem cells (*ISCs*) and their differentiation into enterocytes (*ECs*) via ***egfr***^***1***^ (epidermal growth factor receptor) signalling which are indispensable for maintaining homeostasis and renewing the damaged cells/tissues. Figure created using Biorender.

**Figure S2.**
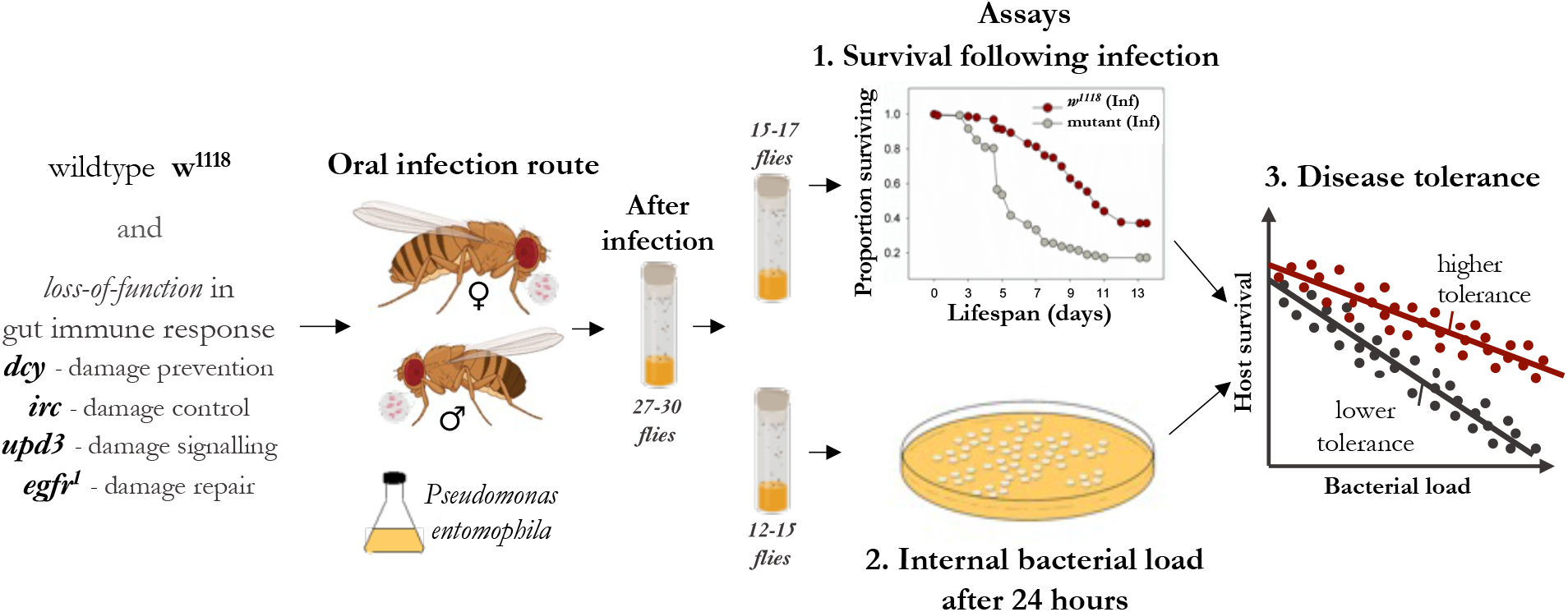
Split-vial experimental design to assay (1) survival (15-17 flies/combination in a vial) and (2) internal bacterial load (12-15 flies/combination in a vial) following oral bacterial infection with Pseudomonas entomophila to test how epithelial immune response including damage prevention and tissue repair mechanisms in Drosophila gut contribute to (3). disease tolerance. For the split-vial design, after oral exposure with each vial containing 25 flies (of each infection treatment, sex, fly line combination) were divided into 2 vials, each for survival (15-17 flies) and bacterial load (12-15 flies). Each point in disease tolerance panel (3) represents replicate-matched data [n=30 vials/infection treatment/sex/fly line – with each vial containing 27-30 flies] from survival (1) and bacterial load (2).

**Figure S3.**
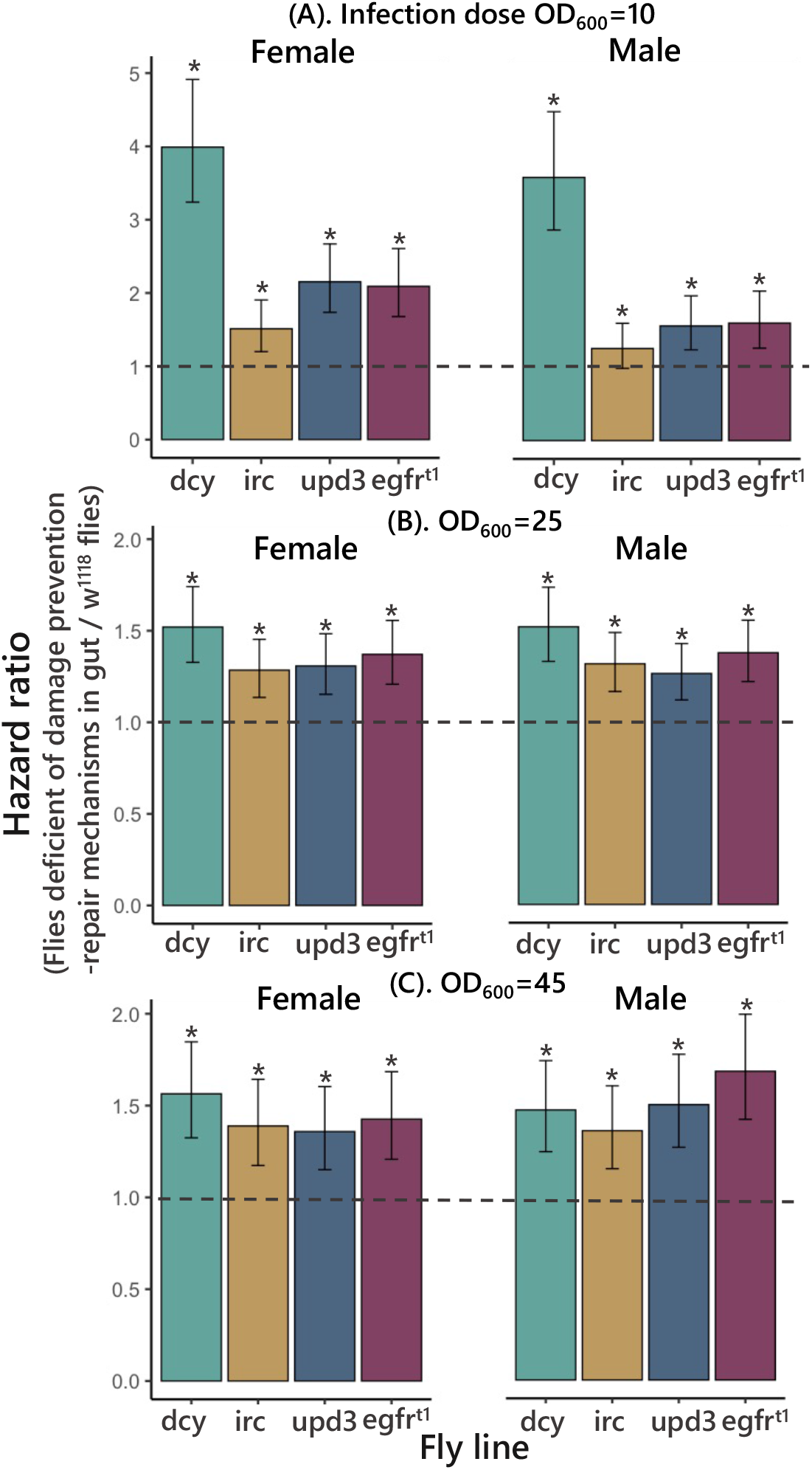
Estimated hazard ratios for males and female flies (*w*^*1118*^ control flies and flies with absence of fully functional gut-epithelial response) calculated from the survival curves. A greater hazard ratio (>1) indicates higher susceptibility to infection. [‘*’ indicates that the transgenic flies are significantly different from *w*^*1118*^ flies for males and females respectively].

**Figure S4.**
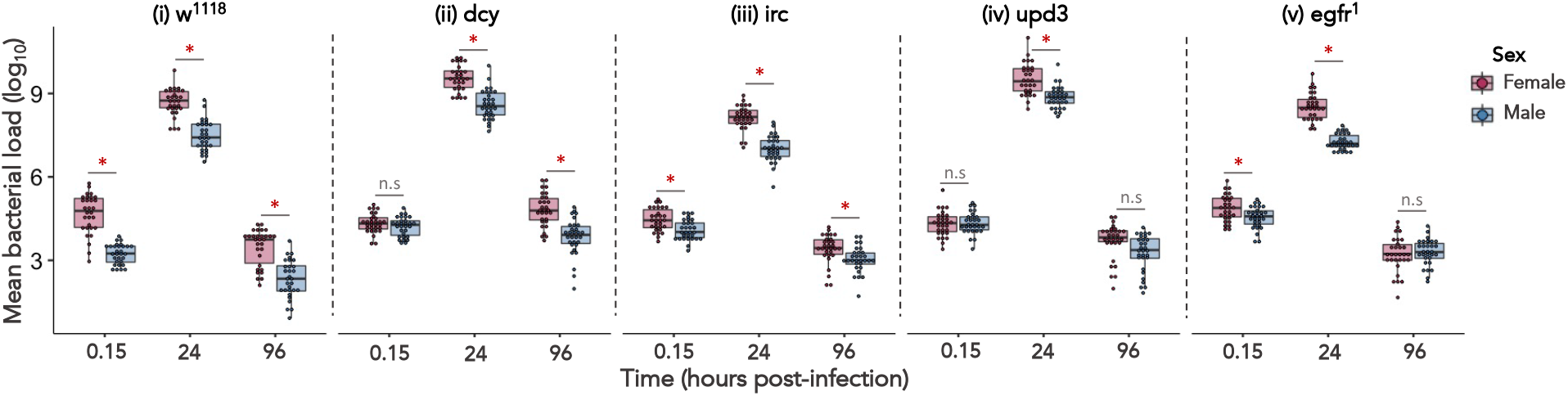
Bacterial load measured at 24 hours after OD600=25 of oral *Pseudomonas entomophila* infection for male & female flies of *w*^*1118*^ and flies with deficient damage repair mechanisms in *Drosophila* gut. “*’ indicates significant differences in males and females of each fly line (w^1118^ and transgenic flies) analysed using Kruskal-Wallis test.

### Supplementary tables

**Table SI-1.**
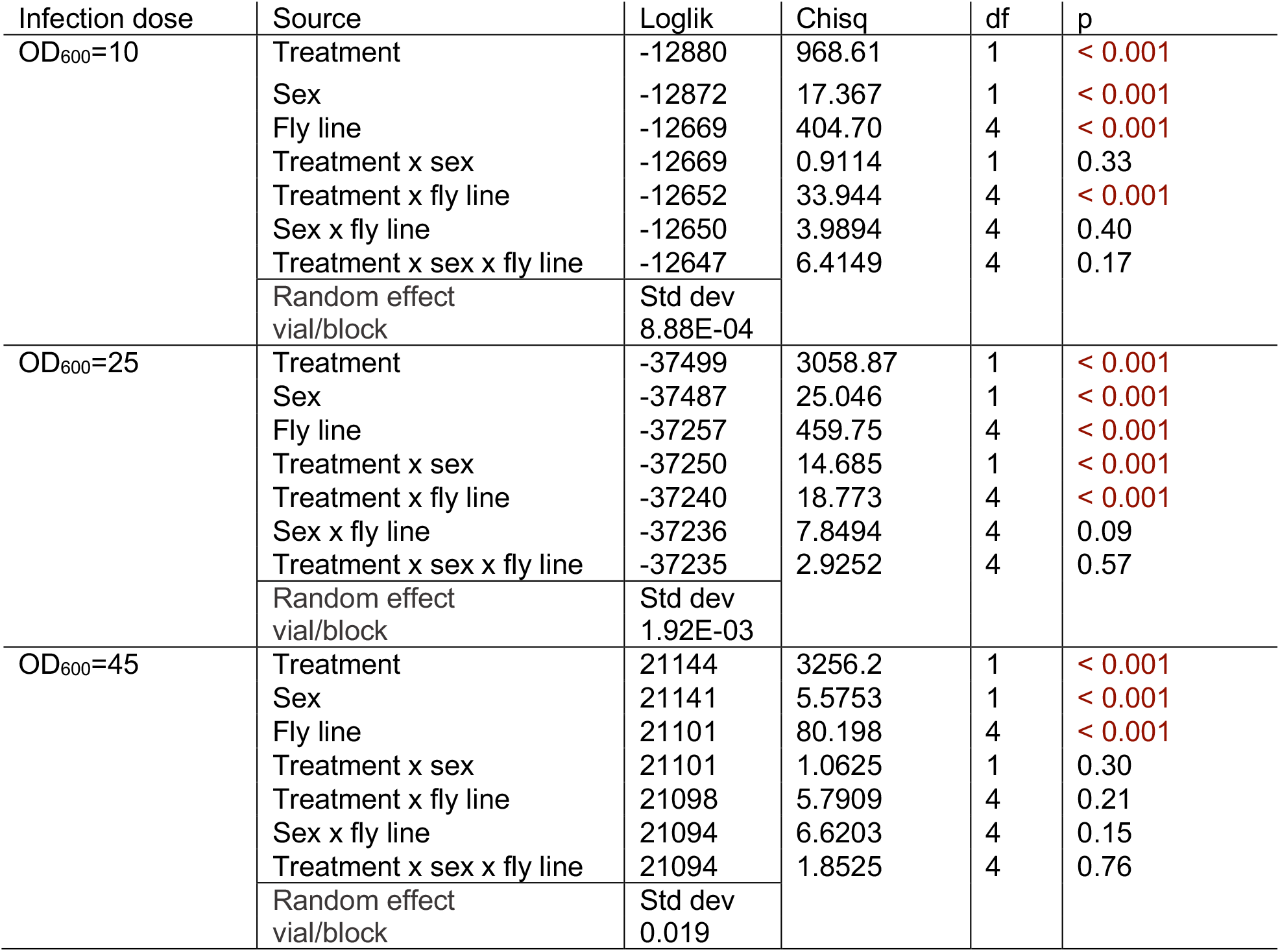
Summary of mixed effects Cox model, for male and female *w*^*1118*^ & flies with *loss-of-function* in in the gut-epithelial response orally infected with 3 different concentration/doses of *P.entomophila*. We used 3-5 day old adult males and females of each fly lines (w^1118^ & flies with deficient damage repair mechanisms in gut) and specified the model as: Survival ∼ Treatment x Sex x Fly line (1|vial/block), with ‘Treatment’, ‘Sex’ & ‘Fly line’ as fixed effects, and ‘vials’ nested in ‘block’ as a random effects. The table shows model output (ANOVA) for survival post oral infection for *w*^*1118*^ flies & flies with *loss-of-function* in in the gut-epithelial response.

| Infection dose | Source | Loglik | Chisq | df | p |
| --- | --- | --- | --- | --- | --- |
| OD<sub>600</sub>=10 | Treatment | -12880 | 968.61 | 1 | < 0.001 |
|  | Sex | -12872 | 17.367 | 1 | < 0.001 |
|  | Fly line | -12669 | 404.70 | 4 | < 0.001 |
|  | Treatment x sex | -12669 | 0.9114 | 1 | 0.33 |
|  | Treatment x fly line | -12652 | 33.944 | 4 | < 0.001 |
|  | Sex x fly line | -12650 | 3.9894 | 4 | 0.40 |
|  | Treatment x sex x fly line | -12647 | 6.4149 | 4 | 0.17 |
|  | Random effect vial/block | Std dev 8.88E-04 |  |  |  |
| OD<sub>600</sub>=25 | Treatment | -37499 | 3058.87 | 1 | < 0.001 |
|  | Sex | -37487 | 25.046 | 1 | < 0.001 |
|  | Fly line | -37257 | 459.75 | 4 | < 0.001 |
|  | Treatment x sex | -37250 | 14.685 | 1 | < 0.001 |
|  | Treatment x fly line | -37240 | 18.773 | 4 | < 0.001 |
|  | Sex x fly line | -37236 | 7.8494 | 4 | 0.09 |
|  | Treatment x sex x fly line | -37235 | 2.9252 | 4 | 0.57 |
|  | Random effect vial/block | Std dev 1.92E-03 |  |  |  |
| OD<sub>600</sub>=45 | Treatment | 21144 | 3256.2 | 1 | < 0.001 |
|  | Sex | 21141 | 5.5753 | 1 | < 0.001 |
|  | Fly line | 21101 | 80.198 | 4 | < 0.001 |
|  | Treatment x sex | 21101 | 1.0625 | 1 | 0.30 |
|  | Treatment x fly line | 21098 | 5.7909 | 4 | 0.21 |
|  | Sex x fly line | 21094 | 6.6203 | 4 | 0.15 |
|  | Treatment x sex x fly line | 21094 | 1.8525 | 4 | 0.76 |
|  | Random effect vial/block | Std dev 0.019 |  |  |  |

**Table SI-2.** Summary of estimated hazard ratio relative to w^1118^ from the cox proportional model. A hazard ratio (>1) indicates that flies lacking damage prevention and repair mechanisms in gut-epithelia are more susceptible to oral *P. entomophila* infection than w^1118^ flies.

|  | Fly line | Sex | Estimate | p | lower 95% | upper 95% |
| --- | --- | --- | --- | --- | --- | --- |
| OD10 | *dcy* | Female | 3.99 | <0.01 | 3.24 | 4.91 |
|  |  | Male | 3.59 | <0.01 | 2.87 | 4.49 |
|  | *Egfr<sup>1</sup>* | Female | 2.09 | <0.01 | 1.68 | 2.61 |
|  |  | Male | 1.60 | <0.01 | 1.26 | 2.04 |
|  | *irc* | Female | 1.51 | 0.0005 | 1.20 | 1.90 |
|  |  | Male | 1.26 | 0.065 | 0.99 | 1.60 |
|  | *upd3* | Female | 2.15 | <0.01 | 1.74 | 2.67 |
|  |  | Male | 1.56 | 0.0002 | 1.24 | 1.98 |
| OD25 | *dcy* | Female | 1.52 | <0.01 | 1.32 | 1.74 |
|  |  | Male | 1.52 | <0.01 | 1.32 | 1.73 |
|  | *Egfr<sup>1</sup>* | Female | 1.37 | <0.01 | 1.20 | 1.55 |
|  |  | Male | 1.37 | <0.01 | 1.21 | 1.55 |
|  | *irc* | Female | 1.28 | <0.01 | 1.13 | 1.45 |
|  |  | Male | 1.31 | <0.01 | 1.16 | 1.48 |
|  | *upd3* | Female | 1.31 | <0.01 | 1.15 | 1.48 |
|  |  | Male | 1.26 | <0.01 | 1.11 | 1.42 |
| OD45 | *dcy* | Female | 1.56 | <0.01 | 1.32 | 1.84 |
|  |  | Male | 1.47 | <0.01 | 1.25 | 1.74 |
|  | *Egfr<sup>1</sup>* | Female | 1.43 | <0.01 | 1.21 | 1.68 |
|  |  | Male | 1.69 | <0.01 | 1.42 | 2.00 |
|  | *irc* | Female | 1.39 | <0.01 | 1.17 | 1.64 |
|  |  | Male | 1.36 | 0.0003 | 1.15 | 1.61 |
|  | *upd3* | Female | 1.36 | 0.0003 | 1.15 | 1.60 |
|  |  | Male | 1.50 | <0.01 | 1.27 | 1.78 |

**Table SI-3.** Summary of log_10_ transformed bacterial load data after exposure to 3 different concentration (doses) *P. entomophila* orally infected flies (w^1118^ & transgenic flies), analysed using non-parametric Kruskal-Wallis test by fitting ‘Sex’ as categorical fixed-effects for timepoints 15 mins, 24 hours & 96 hours for OD_600_=25)

|  | Sex | Fly line |  | Mean diff | Z | p |
| --- | --- | --- | --- | --- | --- | --- |
| 15 min | Female | egfr1 | w<sup>1118</sup> | 3.50 | 0.77 | 0.43 |
|  |  | irc | w<sup>1118</sup> | -7.83 | -1.73 | 0.08 |
|  |  | upd-3 | w<sup>1118</sup> | -11.03 | -2.44 | 0.01 |
|  |  | dcy | w<sup>1118</sup> | -11.16 | -2.47 | 0.01 |
|  | Male | egfr1 | w<sup>1118</sup> | 29.56 | 6.55 | < 0.001 |
|  |  | dcy | w<sup>1118</sup> | 28.96 | 6.423 | < 0.001 |
|  |  | upd-3 | w<sup>1118</sup> | 28.96 | 6.42 | < 0.001 |
|  |  | irc | w<sup>1118</sup> | 28.033 | 6.21 | < 0.001 |
|  | Sex | Fly line |  | Mean diff | Z | p |
| 24 hours load | Female | dcy | w<sup>1118</sup> | 24.30 | 5.388 | < 0.001 |
|  |  | upd-3 | w<sup>1118</sup> | 22.23 | 4.930 | < 0.001 |
|  |  | egfr1 | w<sup>1118</sup> | -7.233 | -1.604 | 0.108 |
|  |  | irc | w<sup>1118</sup> | -18.96 | -4.206 | < 0.001 |
|  | Male | upd-3 | w<sup>1118</sup> | 28.70 | 6.364 | < 0.001 |
|  |  | dcy | w<sup>1118</sup> | 25.90 | 5.743 | < 0.001 |
|  |  | egfr1 | w<sup>1118</sup> | -7.233 | -1.604 | 0.10 |
|  |  | irc | w<sup>1118</sup> | -14.16 | -3.141 | 0.001 |
|  | Sex | Fly line |  | Mean diff | Z | p |
| 96 hours load | Female | dcy | w<sup>1118</sup> | 26.90 | 5.965 | < 0.001 |
|  |  | upd-3 | w<sup>1118</sup> | 6.100 | 1.352 | 0.17 |
|  |  | irc | w<sup>1118</sup> | -5.166 | -1.145 | 0.25 |
|  |  | egfr1 | w<sup>1118</sup> | -8.566 | -1.899 | 0.057 |
|  | Male | dcy | w<sup>1118</sup> | 26.50 | 5.876 | < 0.001 |
|  |  | egfr1 | w<sup>1118</sup> | 22.73 | 5.041 | < 0.001 |
|  |  | upd-3 | w<sup>1118</sup> | 20.36 | 4.516 | < 0.001 |
|  |  | irc | w<sup>1118</sup> | 17.56 | 3.895 | < 0.001 |

